# Insights into polyprotein processing and RNA-protein interactions in foot-and-mouth disease virus genome replication

**DOI:** 10.1101/2023.01.26.525530

**Authors:** Danielle M. Pierce, David J. Rowlands, Nicola J. Stonehouse, Morgan R. Herod

**Affiliations:** School of Molecular and Cellular Biology, Faculty of Biological Sciences and Astbury Centre for Structural Molecular Biology, University of Leeds, Leeds, United Kingdom

**Author notes:** Corresponding author’s.

**Keywords:** FMDV, picornavirus, cleavage, replication, replication complex

## Abstract

Foot-and-mouth disease affects cloven hoofed animals and is caused by foot-and-mouth disease virus (FMDV), a picornavirus with a positive-sense RNA genome. The FMDV genome contains a single open reading frame, which is translated to produce a polyprotein that is cleaved by viral proteases to produce the viral structural and non-structural proteins. Initial processing of the polyprotein occurs at three main junctions to generate four primary products; L^pro^ and the P1, P2 and P3 precursors (also termed 1ABCD, 2BC and 3AB_1,2,3_CD). The 2BC and 3AB_1,2,3_CD precursors undergo subsequent proteolysis to generate non-structural proteins that are required for viral replication, including the enzymes 2C, 3C^pro^ and 3D^pol^. These precursors can be processed through both *cis* and *trans* (i.e., intra- and inter-molecular proteolysis) pathways, which are thought to be important for controlling virus replication. Our previous studies suggested that a single residue in the 3B_3_-3C junction had an important role in controlling 3AB_1,2,3_CD processing. Here, we use *in vitro* based assays to show that a single point mutation at the 3B_3_-3C boundary increases the rate of proteolysis to generate a novel 2C-containing precursor. Complementation assays showed that while this point mutation permitted production of some non-enzymatic non-structural proteins, those with enzymatic functions were inhibited. Interestingly, replication could only be supported by complementation with mutations in *cis* acting RNA elements, providing genetic evidence for a functional interaction between replication enzymes and RNA elements.

**Importance:** Foot-and-mouth disease virus (FMDV) is an economically important pathogen of animals that is responsible for foot-and-mouth disease (FMD). FMD is endemic in many parts of the world and can results in major economic losses. Replication of the virus is a highly coordinated event that occurs within membrane-associated compartments in infected cells and requires the viral non-structural proteins. These are all initially produced as a polyprotein that undergoes proteolysis likely through both *cis* and *trans* pathways (i.e., intra- and inter-molecular proteolysis). Alternative processing pathways can provide a mechanism to help coordinate viral replication by providing temporal control to protein production. Here, we analyse the consequences of mutations that change temporal control of FMDV polyprotein processing. Our data suggests that correct processing is required to produce key enzymes for replication in an environment in which they can interact with essential viral RNA elements. These data further the understanding of FMDV genome replication.

## Introduction

Small RNA viruses have limited genome sequence space and therefore minimal coding capacity. These viruses have evolved several strategies to overcome this limitation, including the use of protein precursors that can perform different functions to the mature proteins. Individual proteins and their precursors can also sometimes perform more than one function (1). Examples are the 3CD and 2BC precursors from poliovirus (PV) and foot-and-mouth disease virus (FMDV), respectively, both members of the *Picornaviridae* family. The 3CD protein is involved in priming genome replication, whilst also having protease activity and the 3C^pro^ and 3D^pol^ cleavage products have protease and RNA-dependent RNA polymerase activities, respectively (1). Likewise, the FMDV 2BC precursor inhibits the secretory pathway, a function apparently independent of the roles of 2B as a viroporin and 2C as a viral ATPase (2–4).

The *Picornaviridae* family includes several important human and animal pathogens, including but not limited to PV and FMDV. PV is responsible for the incapacitating (and potentially fatal) human disease poliomyelitis, while FMDV is the causative agent of foot-and-mouth disease, an acute vesicular disease of cloven-hoofed ruminants including livestock, which can be economically-damaging. The FMDV genome contains a single open reading frame that produces a ~250 kDa polyprotein (5). Initial processing of the FMDV polyprotein occurs at three positions to produce four primary products: L^pro^, the capsid precursor P1-2A and two non-structural protein precursors 2BC (also termed P2) and 3AB_1,2,3_CD (also termed P3) (6, 7). L^pro^ is autocatalytically released from the N-terminal region of the polyprotein (6, 8). The P1-2A precursor is released from the polyprotein via a co-translational 2A-driven ribosome skipping mechanism (9) before the 2BC-3AB_1,2,3_CD polyprotein is processed by 3C^pro^. For FMDV, initial processing is believed to be predominantly at the 2C-3A junction, generating 2BC and 3AB_1,2,3_CD precursors. Further 3C^pro^-mediated proteolysis releases the final proteins via a succession of intermediate precursors (7, 10, 11). Processing of the 2BC precursor ultimately generates the 2B and 2C proteins, both of which have multiple roles in replication.The 3AB_1,2,3_CD precursor is composed of the transmembrane protein 3A, three 3B peptides (individually referred to as 3B_1_, 3B_2_ and 3B_3_,), the protease 3C^pro^, and the polymerase 3D^pol^ (12, 13).

Processing of the non-structural polyprotein by 3C^pro^ is thought to occur through at least two separate pathways to generate mutually-exclusive sets of precursors (14). For example, for FMDV, the 3AB_1,2,3_CD precursor is processed to generate the precursors 3AB_1,2,3_C and 3CD, which must be derived from alternative processing strategies. Likewise, for PV, the 3ABCD precursor (the equivalent of 3AB_1,2,3_CD in FMDV) can be processed to generate 3ABC and 3CD. Furthermore, it appears that this alternative processing may be temporally controlled and used to regulate virus replication. For example, previous studies with PV have demonstrated that later production of 3AB and 3CD can delay the initiation of viral RNA replication (15). For FMDV, reducing cleavage of 3CD inhibits replication by limiting the supply of 3D^pol^ (16, 17). Processing through alternative pathways is likely to be driven (in part) through a switch between intra-molecular vs inter-molecular proteolysis (i.e., *cis* vs *trans* cleavage events). However, methods to differentiate between these *cis* vs *trans* cleavages events are challenging and as a result the mechanism(s) that controls this switch are not completely understood.

Like all positive-sense RNA viruses, picornavirus genome replication is associated with virus-induced cytoplasmic membranous structures, sometimes referred to as “replication complexes” or “replication organelles” (18). In these assemblies, multiple new viral positive-strand RNAs are synthesised via a complementary negative-sense template. For FMDV the full composition of these assemblies is unknown, but they are likely composed of multiple viral and cellular factors, including the non-structural proteins 3B, 3D^pol^ and 3CD. Some of the viral non-structural proteins and precursors associate with RNA elements located in the 5’ and 3’ untranslated regions (UTRs) that flank the open reading frame (5). The 5’ UTR of FMDV is uncharacteristically long for a picornavirus and contains several distinct structural elements, including an internal ribosome entry site (IRES), a *cis*-acting replicative element *(cre)* and a large stem loop (termed the S-fragment) (19–22). The IRES has been well studied and is used to initiate protein translation in a cap-independent manner (20, 21). The *cre* is essential for replication and acts as the template for uridylation of 3B (also known as VPg) to generate the replicative primer, VPg-pUpU (23–28). The role of the S-fragment in FMDV replication is less well understood but may be involved in both replication and modulating the innate immune response (29). In other picornaviruses, such as PV, an RNA structure termed the cloverleaf (or oriL) is located at the 5’ terminus of the genome at the site occupied by the S-fragment stem-loop in FMDV (30). This interacts with the precursor protein 3CD as well as host proteins and is involved in initiating negative-strand RNA synthesis (31–36). Furthermore, for PV, other precursor proteins have also been implicated in 5’ UTR interactions, including 3AB (the equivalent to 3AB_1,2,3_ in FMDV), 3BCD and 3ABCD, and have been suggested to be important for controlling replication (15, 32, 37, 38). The role of precursors in FMDV replication is less well established.

In a previous study, we used an FMDV replicon system, where replication is monitored by fluorescent protein (e.g., GFP/RFP) expression over time, to investigate FMDV replication by mutation of the 3B proteins (16). We reported a series of amino acid substitutions that increased the efficiency of processing at the 3B_3_-3C junction but inhibited replication by abrogating the release of free 3D^pol^. Simultaneously, we observed that these mutations caused an overall shift in 3AB_1,2,3_CD processing and channelled precursor synthesis mainly down one pathway generating 3AB_1,2,3_ and 3CD precursors. This series of mutations has enabled us to separate alternative cleavage pathways and study the function of different precursor sets. Here, we investigated the mechanism by which these mutations result in increased processing between 3B_3_ and 3C. Our data suggest that a single amino acid substitution increases sensitivity to *trans*-mediated proteolysis at this boundary. Furthermore, when placed into the context of a full-length polyprotein, this single substitution resulted in accumulation of a novel precursor. Interestingly, it also prevented reciprocal complementation of replicons *in trans,* which we demonstrate is due to a deficiency in the functions of essential viral enzymes.

## Results

### A single point mutation in 3B_3_ prevents replicon replication

In a previous study, we reported that mutations within 3B_3_, at the boundary with 3C^pro^, dramatically changed processing of the FMDV 3AB_1,2,3C_D polyprotein and prevented release of active 3D^pol^. However, through blind passage a compensatory mutation was selected which restored replication and wild-type (WT) 3AB_1,2,3_CD processing. This was identified as a reversion of a lysine at the P2 residue of the 3B_3_-3C junction to the WT threonine (15). These data suggested that the amino acid in the P2 position of the 3B_3_-3C junction alone can be a major determinant of altered polyprotein processing.

Before investigating the mechanism by which mutations at this junction increased processing, we first sought to establish that this residue alone was sufficient to change FMDV polyprotein processing and prevent replicon replication. To this end, we generated an FMDV replicon with a threonine to lysine mutation at the P2 residue of the 3B_3_-3C cleavage junction (Figure 1A and B). In this replicon (termed, ptGFP-3B_3_^T>K^) the reporter protein ptGFP replaced the structural proteins, allowing ptGFP expression to be used as an indicator of replicon replication (Figure 1A). This replicon RNA was transfected into BHK-21 cells alongside the previously published mutant replicons ptGFP-3B_1,2,3_^Y3F^ (contains inactivating point mutations to the triptych of 3B genes) and ptGFP-3B_3/2_ (the six C-terminal residues of 3B_3_ replaced by those of 3B_2_) (Figure 1B). A WT ptGFP expressing replicon (ptGFP) and a replicon containing an inactivating double point mutation in 3D^pol^ (ptGFP-3D^GNN^) were included as controls. The latter replicon serves as a negative control for ptGFP production from translation of the input transfected RNA, as we have previous described (39). RNAs from these replicons were transfected into BHK-21 cells and replication monitored by ptGFP fluorescence using an Incucyte real-time imaging system (Figure 1C).

**Figure 1.**
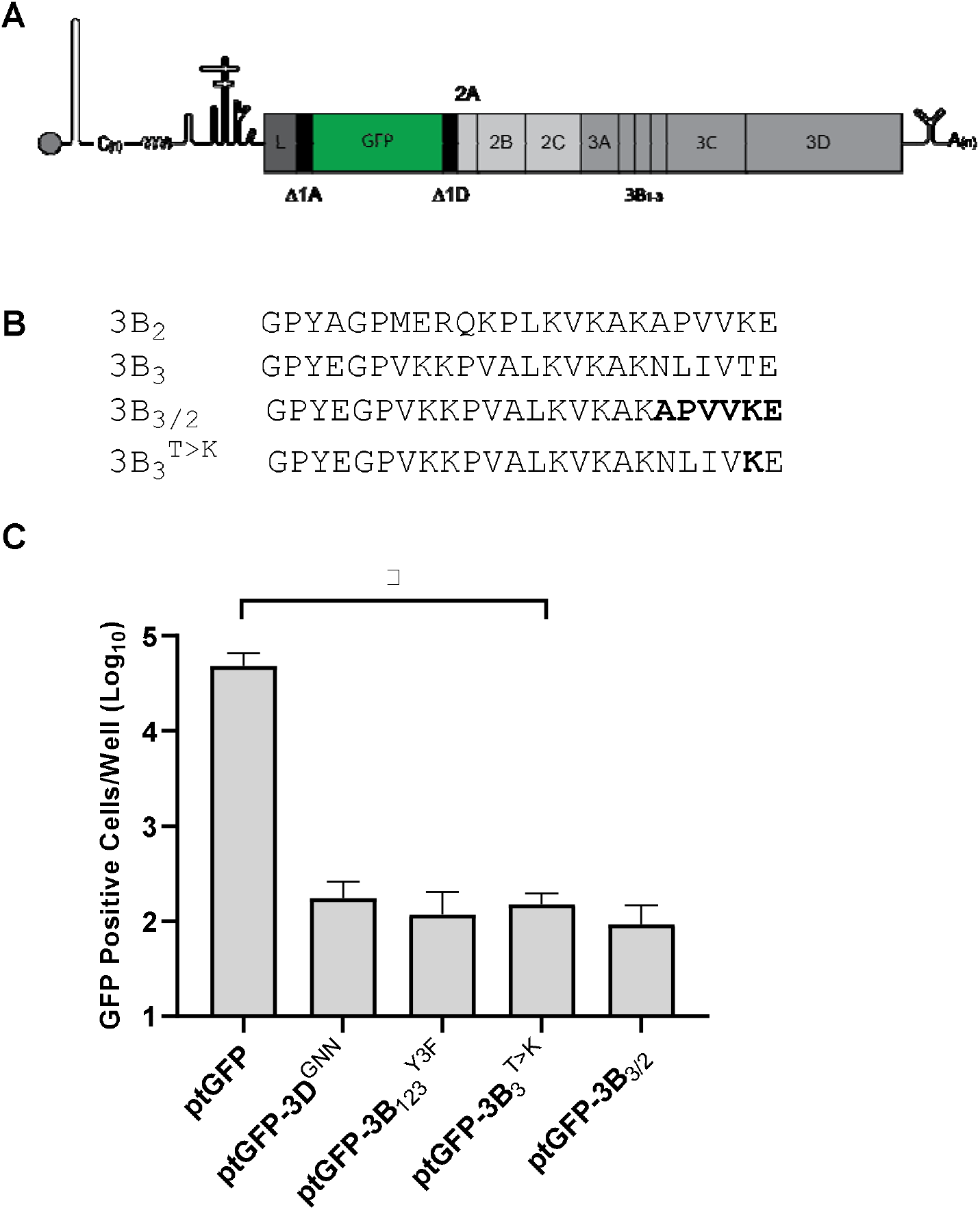
A single mutation at the 3B_3_-3C junction prevents FMDV replicon replication. **(A)** Schematic diagram of the FMDV replicon. **(B)** Sequence alignments of the 3B cleavage junctions with the 3B_3/2_ and 3B_3_^T>K^ mutants. **(C)** Replication of replicons containing 3B_3_ mutations as well as the WT ptGFP replicon (ptGFP) and replication-defective controls containing inactivating mutations in 3D^pol^ (3D^GNN^), or 3B proteins (3B_1,2,3_^Y3F^). GFP expression was monitored hourly for 24 hours. The graph shows GFP positive cells per well at 8 hours post-transfection when replication is maximal. Significance compared to WT control (n = 3 ± SEM; * = p ≤ 0.05).

As anticipated, the WT ptGFP replicon produced ptGFP >100-fold greater than the ptGFP-3D^GNN^ control replicon, as previously reported. In comparison the ptGFP-3B_3_^T>K^ replicon showed GFP expression equivalent to the replication-defective ptGFP-3D^GNN^ control. These data were in agreement with those obtained with the ptGFP-3B_3/2_ and ptGFP-3B_1,2,3_^Y3F^ replication-defective replicons we previously reported (16). Thus, the 3B_3_^T>K^ mutation alone is sufficient to prevent replicon replication.

### A single mutation at the 3B_3_-3C boundary increases the rate of proteolysis

We previously demonstrated that the ptGFP-3B_3/2_ mutation inhibited replication and changed processing of the 3AB_1,2,3_CD polyprotein. To confirm that the 3B_3_^T>K^ substitution was sufficient to induce the same changes, we employed the previously described *in vitro* coupled transcription/translation assay (16). T7 expression constructs were generated to express either the WT FMDV polyprotein or a polyprotein containing the 3B_3_^T>K^ point mutation. The polyprotein used in these experiments included 2BC as well as the 3AB_1,2,3_CD region to determine changes to the entire NS polyprotein. These experiments also included a control polyprotein containing an inactivating mutation in 3C^pro^ (3C^C163A^) predicted to prevent its proteolytic activity (40). Processing was investigated by [^35^S] methionine/cysteine pulse/chase labelling in *in vitro* coupled transcription/translation reactions, harvesting samples at regular time points and analysing protein products by SDS-PAGE (Figure 2A and B).

**Figure 2.**
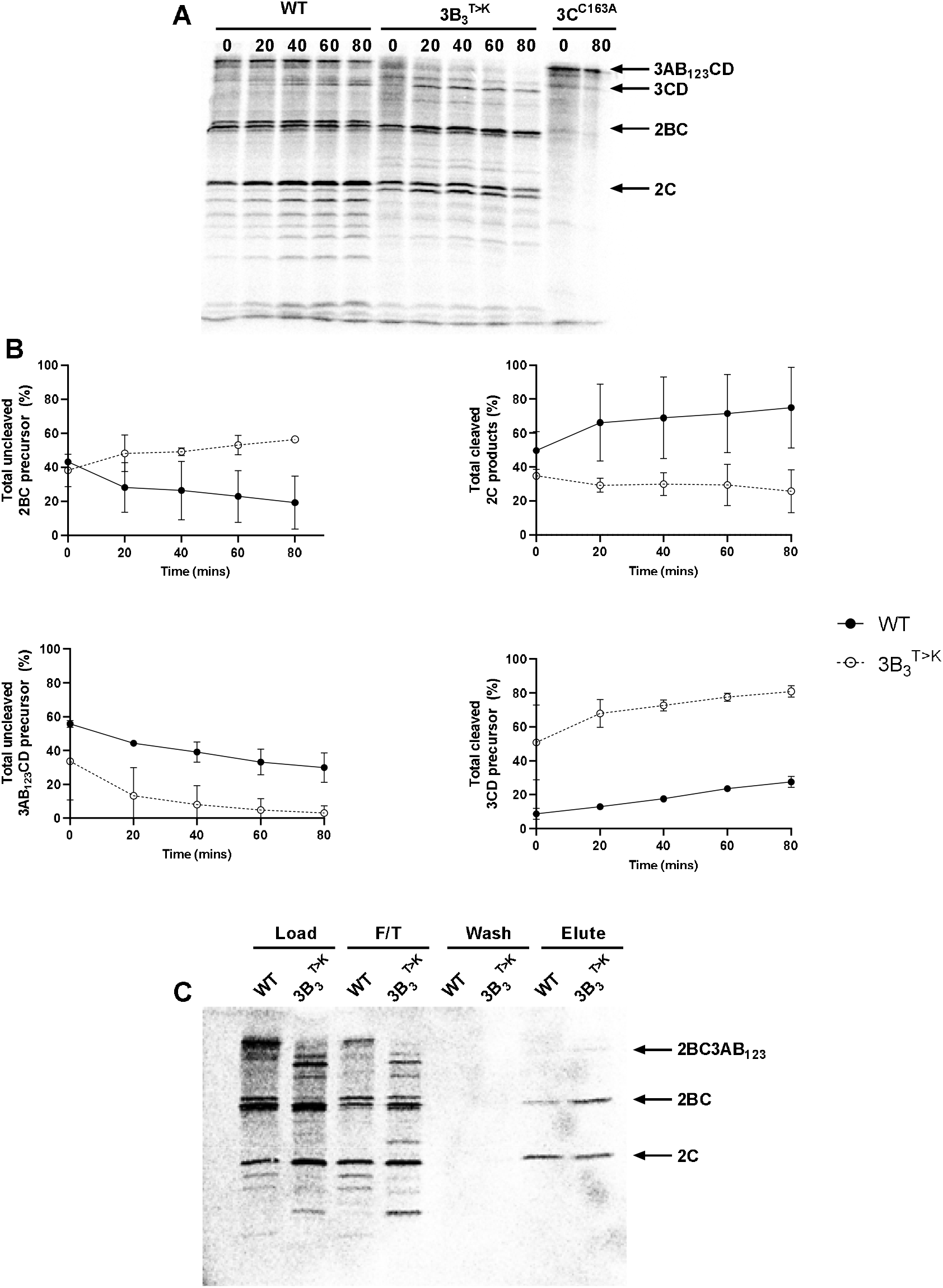
A single mutation at the 3B_3_-3C boundary increases the rate of proteolysis and drives the production of novel precursors. **(A)** Plasmids expressing the WT or mutant 3B_3_^T>K^ FMDV polyprotein precursors were used to prime [^35^S] labelled pulse-chase *in vitro* coupled transcription/translation reactions. At regular intervals samples were taken and stopped by the addition of 2 x Laemmli buffer. Proteins were separated by SDS-PAGE and visualised by autoradiography. The identity of some FMDV proteins is shown. **(B)** The percentage of protein or protein precursor was quantified as a total percentage of 2C or 3D^pol^ containing products, as appropriate (n = 2 ± SD). **(C)** Duplicate reactions were incubated for 90 minutes before immunoprecipitation of 2C containing precursors with anti-2C antibodies. The pre- and post-precipitation samples were separated by SDS-PAGE and visualised by autoradiography. Arrows show the identity of 2C containing proteins, based on predicted molecular weights.

For the WT construct, the full-length 3AB_1,2,3_CD precursor was detected at early time points, and was steadily processed over time primarily into 3AB_1,2,3_C and 3D^pol^, with a small amount of 3CD derived from an alternative processing pathway. At the later time points (40 and 60 minutes), 3AB_1,2,3_ was also detected. Both 2B and 2C were present, in addition to a limited amount of the precursor 2BC at earlier time points. When compared to WT, the construct containing the 3B_3_^T>K^ mutation resulted in greater amounts of the 3CD and 3AB_1,2,3_ precursors and less of the 3AB_1,2,3_CD precursor and 3D^pol^. There were also increased levels of the 2BC precursor in addition to a high molecular weight precursor, possibly 2BC3AB_1,2,3_, which was detected at early time points and gradually decreased over time. These data extended our previous observations demonstrating that the 3B_3_^T>K^ substitution alone is sufficient to accelerate proteolysis at the 3B_3_-3C junction and so increases the relative amounts of 2BC, 3CD and 3AB_1,2,3_.

### Increasing polyprotein proteolysis generates a novel 2BC containing precursor

The previous *in vitro* translation experiments suggested that the 3B_3_^T>K^ mutation increased the production of the 2BC precursor in addition to a larger molecular weight precursor not observed in the WT control. To confirm the identity of the 2C-containing precursors the *in vitro* translation samples were immunoprecipitated using an anti-2C antibody. T7 constructs expressing the WT polyprotein or polyprotein containing the 3B_3_^T>K^ substitution were used in *in vitro* coupled transcription/translation reactions with [^35^S] methionine/cysteine pulse/chase labelling. Samples were taken after 90 minutes and immunoprecipitation performed on half of the sample. Both pre- and post-immunoprecipitation protein samples were analysed by SDS-PAGE (Figure 2C).

In comparison to WT, a smaller proportion of mature 2C (but more unprocessed 2BC precursor) were immunoprecipitated with an anti-2C antibody following expression from the polyprotein containing the 3B_3_^1 K^ mutation. Furthermore, the additional higher molecular weight band which was only present following expression of the 3B_3_^T>K^ precursor was also immunoprecipitated with the anti-2C antibody. Based on these observations and the estimated molecular weight, this product is mostly likely a 2BC3AB_1,2,3_ precursor. These results agree with previous data and suggest that the 3B_3_^T>K^ mutation preferentially increases the rate of proteolysis at the 3B_3_-3C junction, compared to the 2C-3A junction, resulting in the accumulation of 2BC3AB_1,2,3_, which is not normally detected.

### The 3B_3_^T>K^ substitution stimulates *trans*-mediated processing

We speculated that there were two likely mechanisms by which the order of polyprotein processing was altered. It is possible that the point mutation affected protein folding conformations to change the order of *cis*-mediated proteolysis such that the mutant 3B_3_^T>K^ boundary was processed first. However, we believe the more likely possibility is that the point mutation generated a boundary sequence that was preferentially recognised *in trans* by 3C^pro^ and/or a 3C^pro^-containing precursor. To explore the latter possibility, we adapted our *in vitro* assay to investigate *trans*-mediated cleavage. For simplicity, we adapted both the WT 3AB_1,2,3_CD precursor or precursor containing the 3B_3_^T>K^ substitution to also contain the inactivating mutation in 3C^pro^ (3C^C163A^) to prevent self-proteolysis (40). These precursor substrates (termed 3C^C163A^ and 3B_3_^T>K^-3C^C163A^, respectively) were translated *in vitro* with [^35^S] methionine/cysteine before adding excess unlabelled methionine/cysteine and purified active 3C^pro^ to a duplicate set of reactions (plus 3C^pro^). Samples were harvested at regular time points and processing of the [^35^S] labelled precursor was analysed by SDS-PAGE (Figure S1). As controls, the experiment was conducted with the WT and 3B_3_^T>K^ constructs that did not contain the 3C^C163A^ point mutation (Figure S1). To aid interpretation the relative amount of 3AB_1,2,3_CD and 3CD products was quantified by phosphorimaging (Figure 3).

**Figure 3.**
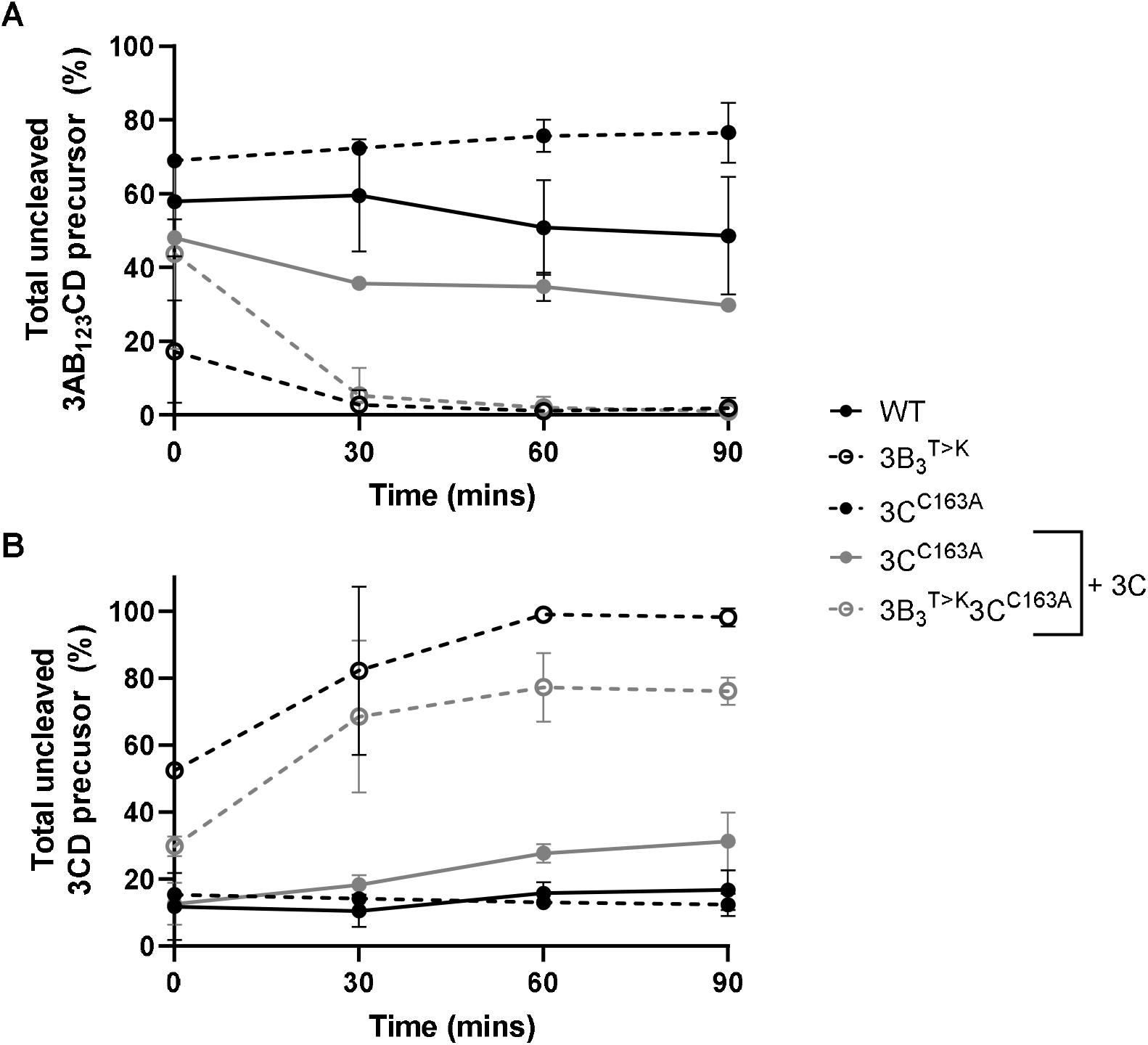
The 3B_3_^1 K^ point mutation drives *trans*-mediated precursor proteolysis. Plasmids expressing proteolytically inactive polyproteins with or without the 3B_3_^T>K^ mutation (termed 3B_3_^T>K^3C^C163A^ and 3C^C163A^, respectively) were used to prime coupled [^35^S] labelled transcription/translation assays, followed by unlabelled amino acid chase. To a duplicate set of reactions 10 μM purified 3C^pro^ was added (+3C) immediately after chase. As controls, reactions were setup alongside with the WT or 3B_3_^T>K^ polyproteins without an inactivated 3C^pro^. At regular intervals, samples were taken and reactions stopped by the addition of 2 x Laemmli buffer. Proteins were separated by SDS-PAGE and visualised by autoradiography. The relative proportion of uncleaved 3AB_1,2,3_CD **(A)** or 3CD **(B)** was quantified as a percentage of 3D^pol^ containing products (n = 2 ± SD).

Both WT and 3B_3_^T>K^ precursors carrying the 3C^C163A^ mutation produced only uncleaved full-length 3AB_1,2,3_CD in the absence of 3C^pro^ provided *in trans,* as anticipated. The addition of 3C^pro^ resulted in the production of smaller proteins due to *trans*-mediated proteolysis of 3AB_1,2,3_CD. For the WT precursor, these were predominantly 3AB_1,2,3_, 3CD and 3D^pol^, in addition to a cluster of 3B_1,2,3_CD precursors, indicative of alternative cleavage pathways. In comparison, the precursor containing the 3B_3_^T>K^ mutation was processed to 3AB_1,2,3_ and 3CD over the duration of the experiment, as observed previously with the active precursor molecule (Figure 2A).

To investigate whether the 3B_3_^T>K^ precursor was also sensitive to cleavage by 3C^pro^ when this is present as part of a larger precursor molecule, we generated a 3AB_1,2,3_CD expression construct in which all the cleavage boundaries had been mutated. Thus, the protease activity of this precursor was retained but only in the context of a full-length 3AB_1,2,3_CD polyprotein. Our *trans*-cleavage assay was repeated using this new construct, termed 3AB_1,2,3_C^pro^D, in place of the purified 3C^pro^ enzyme used above (Figure 4 and Figure S2).

**Figure 4.**
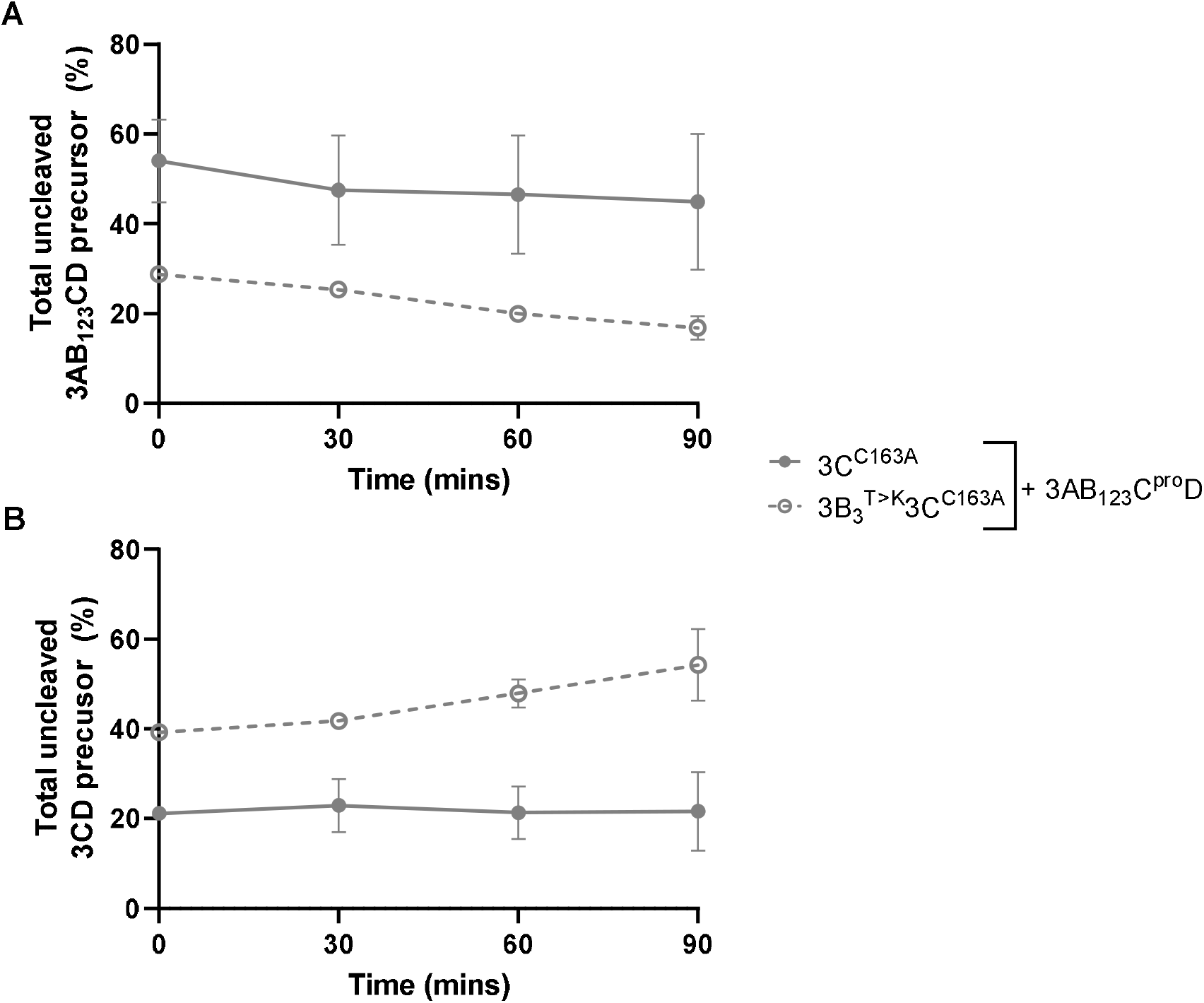
The 3B_3_^T>K^ point mutation drives *trans*-mediated precursor proteolysis. Plasmids expressing proteolytically inactive versions of WT or 3B_3_^T>K^ polyproteins (3C^C163A^ and 3B_3_^T>K^3C^C163A^, respectively) were used to prime coupled [^35^S] labelled transcription/translation assays in the presence of 3AB_1,2,3_C^pro^D, a proteolytically active precursor with all cleavage boundaries mutated to prevent self-proteolysis (+3AB_123_C^pro^D). At regular intervals, samples were taken and reactions stopped by the addition of 2 x Laemmli buffer. Proteins were separated by SDS-PAGE and visualised by autoradiography. The relative proportion of uncleaved 3AB_1,2,3_CD **(A)** or 3CD **(B)** was quantified as a percentage of 3D^pol^ containing products (n = 2 ± SD).

As before, the 3C^C163A^ mutation prevented self-proteolysis when present within the WT precursor or the precursor containing the 3B_3_^T>K^ substitution, as expected. Addition of the proteolytically active 3AB_1,2,3_C^pro^D construct resulted in processing of the proteolytically inactive 3AB_1,2,3_CD precursor bearing the 3B_3_^T>K^ mutation to generate 3AB_1,2,3_ and 3CD. This pattern of processing was similar to that observed following addition of active 3C^pro^, as observed above (Figure 3). This contrasts with the WT proteolytically inactive 3AB_1,2,3_CD precursor, which was not significantly processed *in trans* by 3AB_1,2,3_C^pro^D. Taken together, these data suggest that the 3B_3_^T>K^ mutation at the 3B_3_-3C junction generates a cleavage boundary that is preferentially recognised by 3C^pro^ (even when delivered as part of a larger precursor). Thus, driving rapid *trans*-mediated proteolysis at this junction results in over-production of a specific set of viral precursor proteins.

### Increasing the rate of 3B_3_-3C cleavage prevents production of *trans*-functional 2C and 3D^pol^ but not 3B

The *in vitro* polyprotein processing data above implies that the 3B_3_^T>K^ mutation increases the rate of proteolysis at the 3B_3_-3C junction by stimulating *trans*-mediated proteolysis. In doing so, it drives the formation of 2BC, 3AB_1,2,3_ and 3CD precursors to the detriment of other products such as the enzymes 2C and 3D^pol^. In our previous studies we used *trans*-complementation assays to investigate protein function. These assays involve the co-transfection of two replication defective replicon constructs that express different fluorescent reporter genes, allowing their replication to be differentially monitored. Co-transfection of the two replicons allows exchange of viral non-structural proteins within replication complexes to permit replication of one (or both) of the input genomes (41) (Figure 5A). Here, we used this approach to investigate whether stimulating *trans*-mediated proteolysis at the 3B_3_-3C junction prevented the production of functional 2C. To do this, mutants introduced into 2C were investigated to determine if these could be compensated by a replicon harbouring a 3B_3_^T>K^ mutation. If this is possible, it would indicate that changing the temporal order of polyprotein processing does not prevent 2C function.

**Figure 5.**
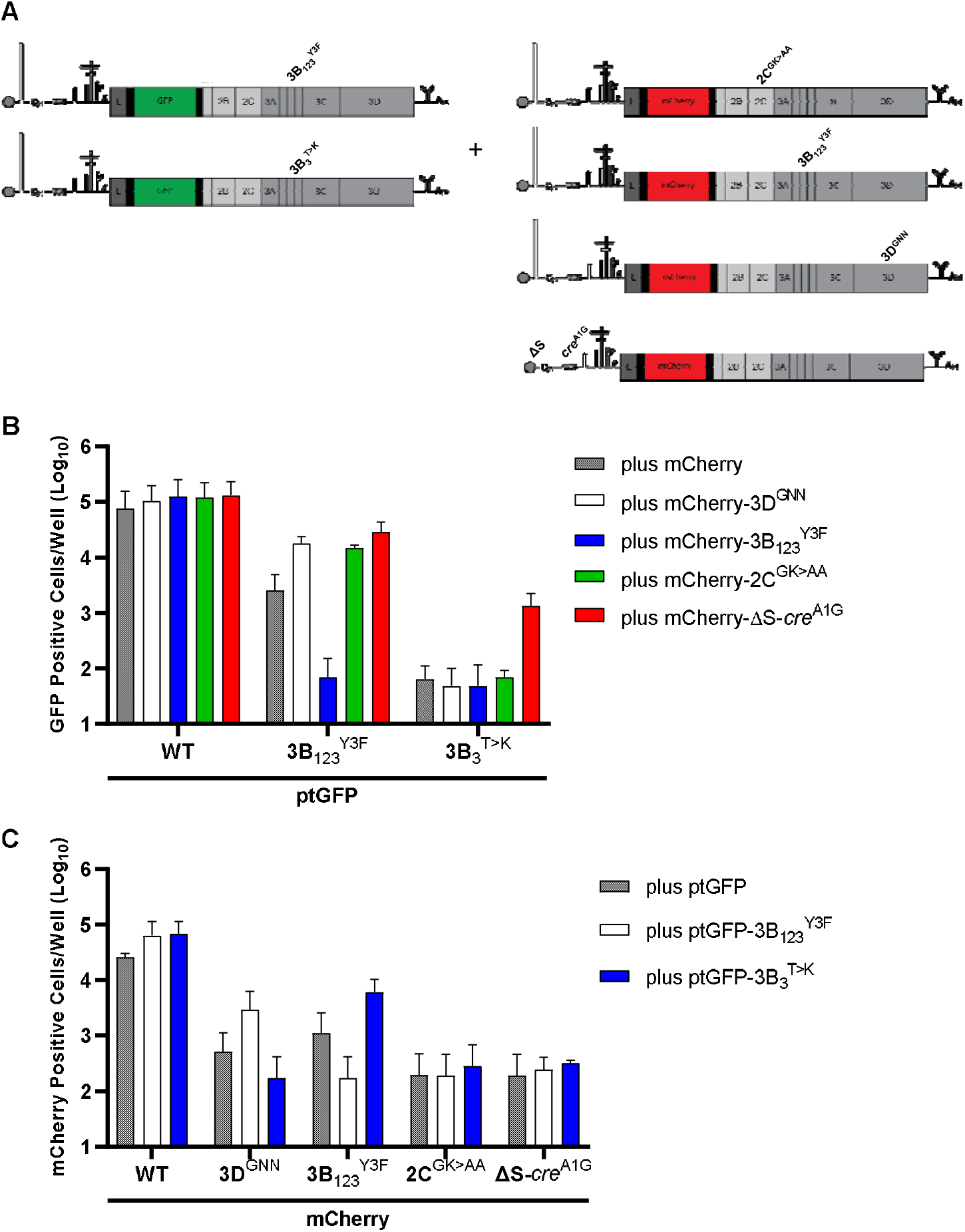
The 3B_3_^T>K^ substitution prevents complementation of 2C mutants *in trans.* **(A)** Schematic of the *trans*-complementation experiment which involved co-transfecting BHK-21 cells with mCherry replicons containing replication-defective 2C or 3B mutations together with a WT ptGFP, ptGFP-3B_3_^T>K^ or ptGFP-3D^GNN^ replicon. Fluorescent protein expression was monitored hourly for 24 hours. The data show **(B)** ptGFP positive cells per well or **(C)** mCherry positive cells per well at 8 hours post-transfection (n = 2 ± SD).

To this end, replication-defective mCherry constructs were generated which contained inactivating mutations at catalytic 2C residues (termed, mCherry-2C^GK^>AA) (42). This construct was co-transfected with the ptGFP-3B_3_^T>K^ replicon (as used in Figure 1C), or a WT ptGFP replicon control. As a positive control, co-transfections were also performed with an mCherry-3B_1,2,3_^Y3F^ replicon (which contains inactivating point mutations to the triptych of 3B genes), which we have shown can be complemented *in trans* (16). Co-transfections were also performed with WT mCherry or ptGFP replicons to eliminate the possibility of any dominant-negative effects and yeast tRNA to act as a negative control for no complementation (Figure 5A). Replication was monitored by both ptGFP and mCherry expression and the number of fluorescent positive cells quantified at 8 hours post-transfection, as documented previously (41). For brevity the key data sets and controls are shown (Figure 5B and 5C) with the complete data set shown in supplementary material (Figure S3).

Replication of the WT mCherry or ptGFP replicon did not significantly change upon co-transfection with any of the RNAs tested, suggestive of no dominant negative effects. Replication of the mCherry-3B_1,2,3_^Y3F^ replicon was significantly enhanced by the ptGFP-3B_3_^T>K^ construct, as anticipated. The mCherry-2C^GK^>AA replicon was not recovered by any of the helper replicons, suggesting the functions of 2C cannot be provided *in trans* (Figure 5C). We also noticed that no functional complementation was provided to the ptGFP-3B_3_^T>K^ mutant replicon by co-transfection with mCherry-3B_1,2,3_^Y3F^. To investigate this further we extended our complementation experiments to include mCherry replicons containing mutations to *cis*-acting RNA replication elements such as the S-fragment or *cre.* The rationale here was that previous studies have demonstrated that deleting *cis*-acting replication elements can improve or allow recovery of replicons *in trans* (41), presumably by increasing the free pool of proteins which would otherwise be sequestered by *cis* interactions. When the ptGFP-3B_3_^T>K^ replicon was co-transfected with a mCherry-ΔS or mCherry-*cre*^A1G^ replicon we observed a significance increase in ptGFP expression, indicating that inactivation of these *cis*-acting replication elements permitted complementation of the ptGFP-3B_3_^T>K^ replicon in *trans* (Figure 5B). Together this data suggests that the ptGFP-3B_3_^T>K^ replicon can supply material *in trans* to recover replicons defective in 3B but not 2C or 3D^pol^ and can only receive complementation *in trans* from replicons with inactivating mutations or deletions to *cis*-acting RNA elements.

### Complementation of replicons with the 3B_3_^T>K^ substitution reveals a requirement for the full-length polyprotein

Having shown that deletion of *cis*-acting RNA elements allowed complementation of the ptGFP-3B_3_^T>K^ replicon, we took advantage of this system to identify which protein component was required for complementation. To this end, we generated a new panel of mCherry replicons in which both the S-fragment and *cre* were inactivated but which expressed just a subset of the polyprotein, namely 3AB_1,2,3_CD, 3B_1,2,3_CD, 3CD or 3D^pol^ (Figure 6A). With these constructs, we were able to probe whether the ptGFP-3B_3_^T>K^ replicon is missing one of these protein components to initiate replication. This new panel of replicons was used in complementation assays with the same controls as described above, with replication monitored by both ptGFP and mCherry expression, and the number of fluorescent positive cells quantified at 8 hours post-transfection. Again, for clarity the key data sets and controls are shown (Figure 6B and 6C) with the complete data set shown in supplementary material (Figure S4).

**Figure 6.**
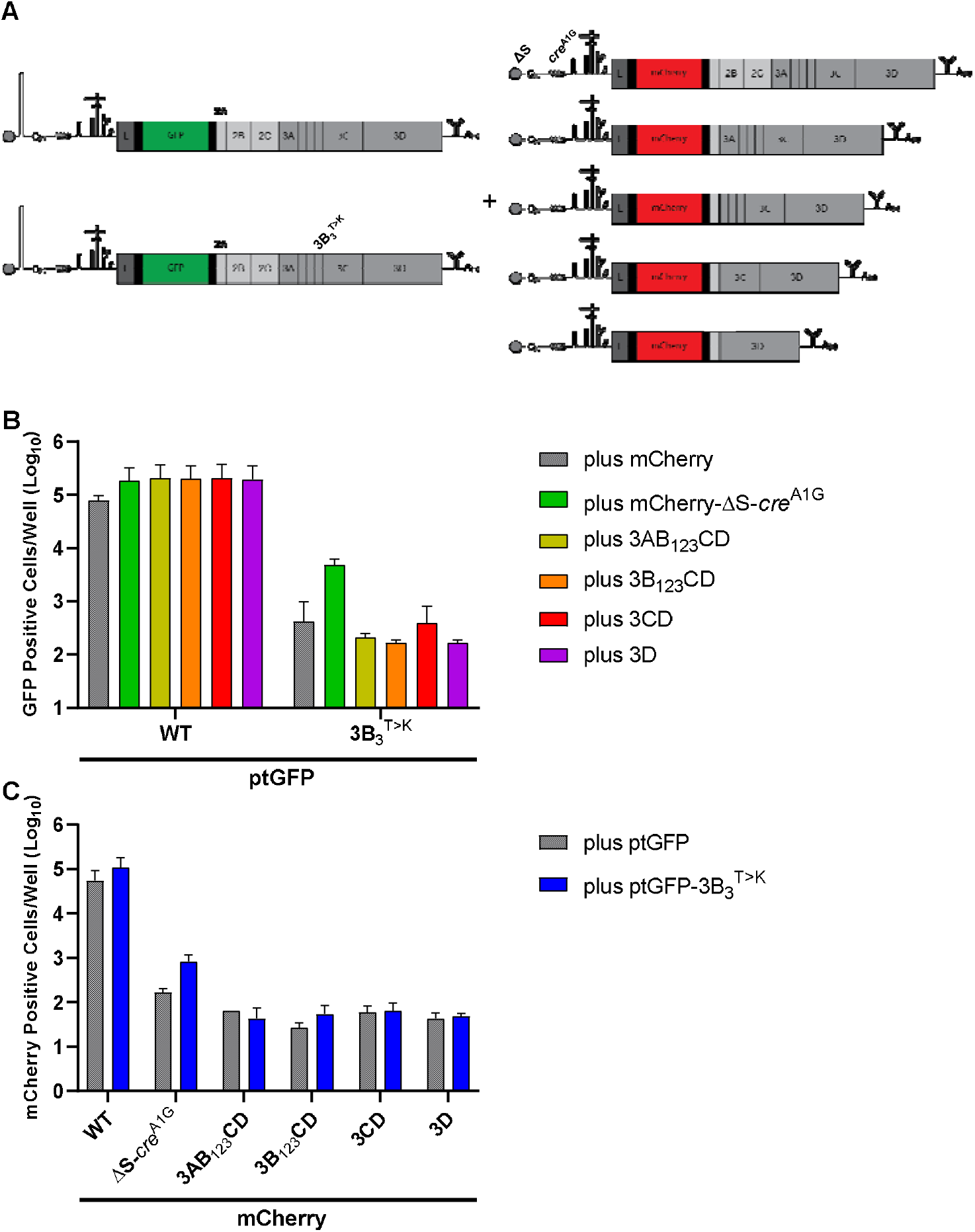
S-fragment deletions allow *trans*-complementation of *cis*-acting replication components. **(A)** Schematic of the trans-complementation experiment which involved co-transfecting BHK-21 cells with mCherry replicons containing S-fragment deletions together with a WT ptGFP, ptGFP-3B_3_^T>K^ or ptGFP-3D^GNN^ replicon. Fluorescent protein expression was monitored hourly for 24 hours. The data show **(B)** ptGFP positive cells per well or **(C)** mCherry positive cells per well at 8 hours post-transfection (n = 2 ± SD).

As described above we found that replicons lacking a functional S-fragment and/or *cre*, were able to significantly increase the replication of the ptGFP-3B_3_^T>K^ replicon, with ptGFP expression increasing >10-fold compared to the controls. In contrast, no complementation of the ptGFP-3B_3_^T>K^ replicon was observed when co-transfected with RNAs expressing just 3AB_1,2,3_CD, 3B_1,2,3_CD, 3CD or 3D^pol^. Hence it would appear that 3AB_1,2,3_CD is not sufficient to support replication of a 3B_3_^T>K^ replicon and provision of 2BC containing proteins is also required.

### The 3B_3_^T>K^ mutation does not prevent RNA-protein interactions

A key observation from our *trans*-complementation work is that the 3B_3_^T>K^ mutation prevents complementation of this non-functional replicon unless the *cis*-acting S-fragment or *cre* elements are deleted from the helper RNA. A possible explanation is that the altered cleavage pattern induced by the 3B_3_^T>K^ mutation prevents replication components (e.g., the 2C helicase or 3D^pol^) from interacting with the template RNA, thus preventing its replication.

To investigate this possibility, we adapted a proximity ligation assay (PLA) to study interactions between replicon RNA and components of the replication complex. Firstly, we *in vitro* transcribed ptGFP-3B_3_^T>K^ replicon RNA (or ptGFP-3D^GNN^ and ptGFP-3B_1,2,3_^Y3F^ controls) in the presence of BrUTP to generate BrU labelled replicon RNA. Alongside, a WT ptGFP was transcribed with BrUTP and used to confirm that BrU labelling had no significant inhibitory effect on replicon replication (data unincluded). The ptGFP-3B_3_^T>K^ BrU replicon was co-transfected into BHK-21 cells seeded on coverslips together with a WT replicon in which 3D^pol^ had been labelled with a HA epitope (termed 3D^HA^). We have previously demonstrated that tagging 3D^pol^ with HA in this manner does not affect replication (41). The coverslips were fixed and the interaction between the BrU-3B_3_^T>K^ RNA and 3D^HA^ probed by PLA using anti-BrU and anti-HA antibodies. This allows RNA-protein interactions between the ptGFP-BrU-3B_3_^T>K^ RNA and 3D^HA^ protein to be measured *in situ.* A positive PLA signal indicates that the mutant replicon RNA can associate with enzymes of the replication complex provided *in trans.* No signal indicates a lack of association (Figure 7).

**Figure 7.**
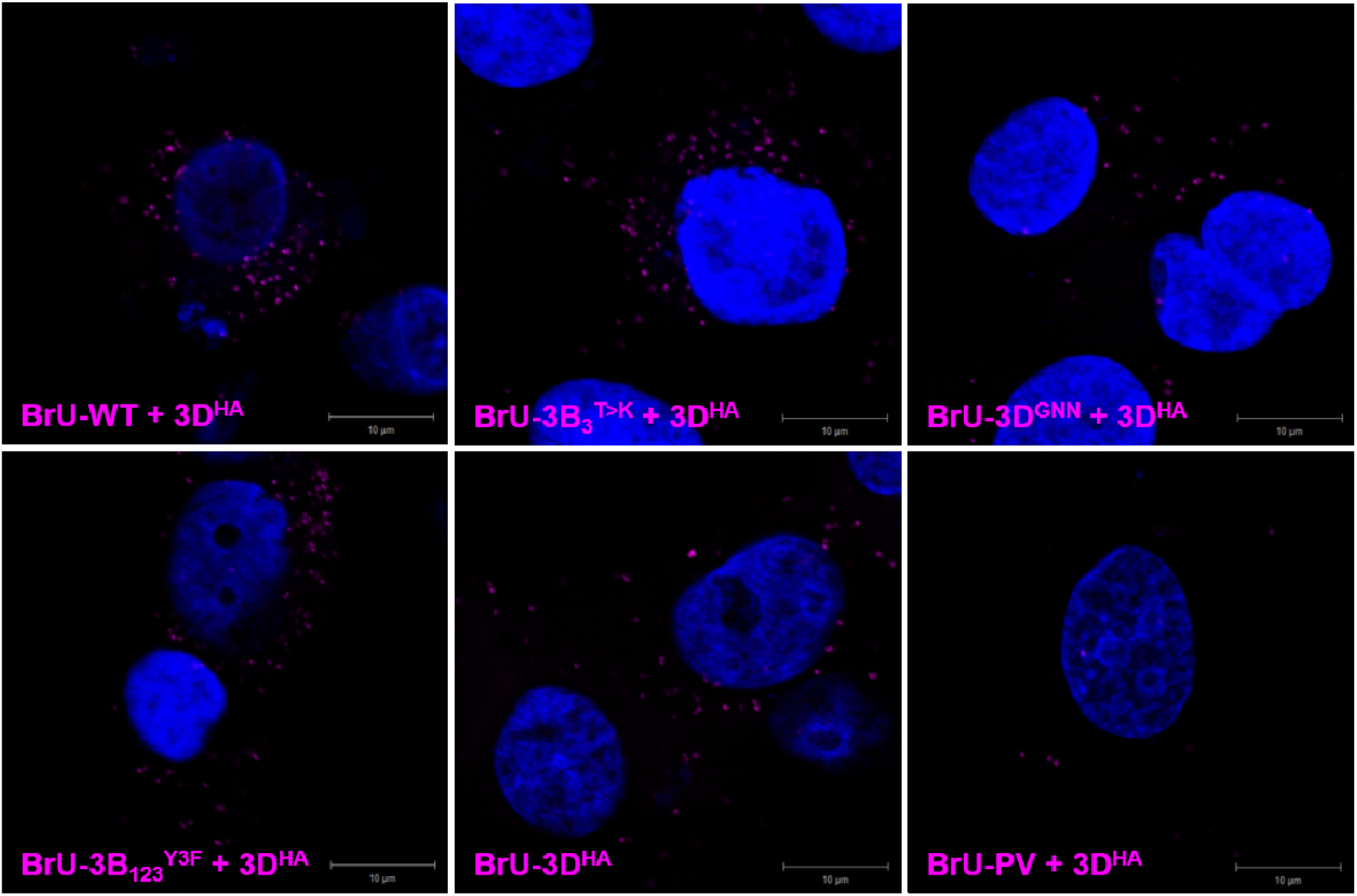
Detection of FMDV RNA-protein complexes by PLA. BHK-21 cells were co-transfected with BrU-labelled ptGFP replicon RNA together with 3D^HA^-labelled replicon RNA. At 4 hours post-transfection cells were fixed and 3D^HA^-BrU RNA complexes detected by proximity ligation assay (PLA) using anti-HA and anti-BrU primary antibodies together with PLA-labelled secondary antibodies. The *in-situ* PLA signal is detected as foci in the cell cytoplasm (pseudo-coloured magenta). Cell nuclei were stained with DAPI (pseudo-coloured blue). Images were captured on a Zeiss LSM-880 confocal microscope (bar 10 μm).

Co-transfection of the WT ptGFP BrU labelled replicon with the WT 3D^HA^ replicon generated PLA signals that were easily detectable, indicative of interactions between the ptGFP RNA and 3D^HA^ protein as predicted from mixing of replication complex components. In contrast, little or no PLA signal was detected when a WT BrU labelled poliovirus replicon (BrU-PV) was co-transfected with the 3D^HA^ helper replicon, suggesting the PLA signal is generated through specific mixing of replication complexes. Co-transfection of the ptGFP-3B_3_^T>K^, ptGFP-3B_1,2,3_^Y3F^ or ptGFP-3D^GNN^ BrU labelled replicon RNAs with the WT 3D^HA^ replicon also generated PLA signals that were easily detectable, albeit at a lower level than in the control sample. This would suggest the 3B_3_^T>K^ mutation does not prevent recruitment of its cognate RNA with a functional RNA polymerase.

## Discussion

All well-studied positive-sense RNA viruses produce polyproteins that help compensate for a relatively limited coding capacity. These polyproteins are processed by viral protease(s) to generate mature proteins via functional intermediates in a highly regulated manner that modulates viral replication. However, establishing how polyprotein processing regulates viral replication can be challenging due to the intricate nature of these interactions. In a previous study of the FMDV polyprotein we showed that changing six amino acids at the 3B_3_-3C cleavage junction prevents viral replication by disrupting polyprotein processing (16). We postulated that this observation provided an opportunity to investigate the role of FMDV polyproteins in viral genome replication. First we investigated whether consequences of substituting the six amino acids could be replicated by a single change. To do this we targeted the P2 residue at the 3B_3_-3C cleavage junction, which from our previous mutational investigation appeared to be the most important for dictating cleavage efficiency. A single lysine substitution was introduced in this position, which we predicted would increase processing and prevent replication. Using a GFP encoding replicon, we showed this single substitution (3B_3_^T>K^) prevented viral replication and in *in vitro* translation assays changed polyprotein processing, as predicted.

To build a more complete picture of how the 3B_3_^T>K^ construct changed polyprotein processing and understand the mechanism that underpinned this change, we employed a combination of *in vitro* translation assays, immunoprecipitation and *trans*-cleavage assays in the context of the FMDV non-structural polyprotein. We found that a 3B_3_-3C cleavage junction bearing this mutation was more efficiently cleaved by 3C^pro^ *in trans* compared to WT. Furthermore, the 3B_3_^T>K^ substitution also rendered this cleavage junction sensitive to proteolysis by 3C^pro^ when the protease was present as part of a larger precursor (i.e., as part of another molecule of 3AB_1,2,3_CD). Thus, the consequence of introducing the 3B_3_^T>K^ substitution was to generate a substrate that was more efficiently cleaved (potentially equal to or greater than either 2B-2C or 2C-3A cleavage sites). This changes the subset of proteins produced; generating 2BC3AB_1,2,3_ (that is not typically observed during infection (7, 43)) significantly increasing levels of 3CD and 2BC and reducing levels of the 3AB_1,2,3_CD. The identity of these products was based on both molecular weight and immunoprecipitation experiments. The 3B_3_^T>K^ mutant construct did generate fully-cleaved 2B and 2C at a reduced rate demonstrating that 3C^pro^ as part of 3CD is proteolytically active, but processing of 3AB_1,2,3_ and 3CD was severely impaired as observed previously (16).

Our data are consistent with biochemical investigations using purified 3C^pro^ which showed that charged residues in the P2 position of the cleavage junction are more efficiently recognised (40, 44–46). It is also consistent with the suggestion that 3C^pro^ retains activity as part of a larger precursor but potentially at lower efficiencies (47–50). Our data therefore agrees with a mechanism of polyprotein processing (suggested from studies with other picornaviruses) in which 3C^pro^ as part of a larger precursor can process the WT 2B-2C and/or 2C-3A junctions most efficiently *in trans,* hence providing a level of regulation to the processing cascade. After these initial cleavage events, 3AB_1,2,3_CD is processed more slowly to allow intermediate and mature cleavage products to fulfil their roles in viral replication (37). Processing of the 3AB_1,2,3_CD precursor can thus proceed potentially through both *cis* and *trans* mechanisms (i.e., intra- and inter-molecular cleavage) and can give rise to alternative precursors, for example, 3AB_1,2,3_C and 3CD. It is clearly suggested from our data that *trans* cleavage of one 3AB_1,2,3_CD molecule by another polyprotein is possible. However, the WT precursor was not processed efficiently by 3AB_1,2,3_CD *in trans,* and the observation that 3C^pro^ in the context of 3CD is proteolytically active, but only able to cleave efficiently at 2B-2C and 2C-3A junctions could suggest that other factors (e.g., a *cis* mediated mechanism), are also involved. Picornavirus 3C^pro^ proteins are also implicated in RNA binding and lipid biogenesis (potentially as part of a precursor), and therefore may act as co-factors to dictate processing pathways (30, 51, 52).

Using *trans*-complementation assays we investigated the function of these different sets of precursors in FMDV replication. A replicon containing the 3B_3_^T>K^ substitution (i.e., producing increased level of 2BC, 3AB_1,2,3_ and 3CD), was able to complement defective mutations in the 3B proteins but not 2C or 3D^pol^, suggesting that this replicon can produce active primers for replication, 3B_1_, 3B_2_, and/or 3B_3_ but inactive replication enzymes, 2C^pro^ and 3D^pol^. Interestingly the 3B_3_^T>K^ replicon was only complemented by a replicon lacking *cis*-acting RNA replication elements *(cre,* S-fragment or both) and when the protein components were provided as part of an entire polyprotein. One interpretation of these pieces of data is that a functional interaction between the non-structural polyprotein and viral RNA elements is required to generate functional enzymes for replication (e.g., 2C^pro^ and 3D^pol^ but not 3B). Hence, if processing of the precursor occurs too rapidly it cannot associate with these RNA elements required for replication and could provide a level of temporal control. A similar mechanism has been suggested for PV where two molecules of 3ABCD are required for replication, one which produces 3CD that interacts with viral RNA structures while one produces enzymatically active 3D^pol^ (33, 37, 53, 54). This two molecule model of processing of 3ABCD required for replication would also be compatible with data that suggests that a larger precursor (such as 3BC) is required to deliver 3B for RNA replication (15, 55, 56). This work extends the growing body of evidence suggesting that processing intermediates are essential for controlling temporally and structurally the organisation of the picornavirus replication organelle.

## Materials and Methods

### Cell lines and plasmids

BHK-21 cells obtained from the ATCC (LGC Standard) were maintained in Dulbecco’s modified Eagle’s medium with glutamine (Sigma-Aldrich) supplemented with 10 % FCS, 50 U / mL penicillin and 50 μg / mL streptomycin.

Plasmids carrying wild-type FMDV replicons, pRep-mCherry and pRep-ptGFP, have already been described (39, 57), along with equivalent plasmids containing 3D^GNN^, 3B_3/2_ and ΔS-fragment mutations (16, 39, 41). Mutations within these plasmids were performed by standard two-step overlapping PCR mutagenesis. For coupled *in vitro* transcription and translation experiments pcDNA3.1(+) based expression plasmids were generated by PCR. Briefly, the relevant FMDV sequence was amplified to including flanking *NotI* restriction enzymes and upstream Kozak modified translational start site. The *Not*I digested PCR products were cloned into *Not*I digested pcDNA3.1(+) (Thermo Fisher Scientific). The sequence of all plasmids used in this study was confirmed by Sanger sequencing. The sequences of all primers and plasmids are available on request.

### Coupled transcription and translation reactions

Coupled *in vitro* transcription and translation assays were performed using the TNT Quick Coupled Transcription/Translation system (Promega) as described previously (16). Reactions contained 10 μL lysate with 250 ng of pcDNA T7 expression plasmid and 0.5 μL [^35^S] methionine/cysteine (PerkinElmer). Reactions were incubated at 30°C for 40 minutes chasing with 2 μL of 50 mg / mL unlabelled methionine/cysteine. Reactions were stopped at 20 minute or hourly intervals by the addition of 2 x Laemmli buffer. Samples were separated by SDS-PAGE before visualisation of radiolabelled products by autoradiography.

For the *trans*-cleavage assays, the TNT reactions were supplemented with purified FMDV 3C^pro^ (a kind gift from Dr Tobias Tuthill (58)) to the indicated final concentration from dilution of a 1 mM stock, simultaneous to the addition of unlabelled methionine/cysteine. Reactions were stopped at 20 minute or hourly intervals by the addition of 2 x Laemmli buffer and the 3C^pro^-mediated proteolysis of radiolabelled precursor monitored by SDS-PAGE.

### *In vitro* transcription

Plasmids containing cDNA copies of FMDV replicons were linearised with *Asc*I before being used to generate T7 *in vitro* transcribed RNA as previously described (39, 41). The reaction was incubated at 32°C for 4 hours before being treated with DNase for 20 minutes at 37°C then purified using an RNA clean and concentrate kit (Zymo Research). The RNA quality was checked using a MOPS/formaldehyde agarose gel electrophoresis.

### Replication and complementation assays

BHK-21 cells were seeded into 24-well tissue cultures vessels, allowed to adhere overnight for 16 hours, before duplicate wells were transfected with 1 μg of each *in vitro* transcribed RNA using Lipofectin (Thermo Fisher Scientific) as previously described (39). For co-transfection complementation assays 500 ng of each RNA molecules were mixed prior to the addition of Lipofectin reagent as previously described (41). Fluorescent reporter expression was monitored using an IncuCyte Zoom Dual Colour FLR (Essen BioSciences) live-cell imaging system housed within a humidified incubator scanning hourly up to 24 hours post-transfection. Images were captured and analysed using the associated software for fluorescent protein expression, as previously described (41). Control transfections (untransfected and the 3D^GNN^ transfection for input translation) were used to determine fluorescent thresholds and identify positive objects from background fluorescence. A positive object was determined as having an average fluorescent intensity of >8 green calibration units (GCU; an arbitrary fluorescent unit) and >2 RCU (red calibration units), which were kept constant throughout the experiments. The number of positive cells per well was determined from the average of up to nine non-overlapping images per well. Unless stated otherwise, data are presented as mean fluorescent positive cells per well at 8 hours post-transfection when replication was approximately maximal. For each experiment, the data were analysed as both fluorescent cell counts per well and total fluorescent intensity per well. There was no difference observed when the data were analysed in either way. Unless otherwise stated, statistical analysis was performed using a two-tailed unpaired t-test.

### Immunofluorescence and proximity ligation assays (PLA)

BHK-21 cells seeded onto coverslips were co-transfected with replicon RNA before fixing in 4 % paraformaldehyde and washing with PBS. Immunofluorescence was conducted as previously described (41). Primary antibodies used were sheep anti-BrdU (Sigma-Aldrich), rabbit anti-FMDV 3D (a kind gift from Francisco Sobrino) and mouse anti-HA (Sigma-Aldrich). Proximity ligation assays (PLA) were conducted using the Duolink® In Situ Red Kit (Sigma-Aldrich), following manufacturer’s instructions.

### Immunoprecipitation

Immunoprecipitation reactions were performed using Dynabeads Protein G (Invitrogen). To bind the antibody to magnetic beads, 5 μL of the FMDV 2C antibody (a kind gift from Francisco Sobrino) was mixed with 195 μL PBS and incubated at room temperature with 50 μL magnetic beads, shaking for 1 hour, after which the supernatant was removed from the beads. Transcription and translation reaction samples were mixed with 200 μL PBS and incubated shaking at room temperature with 25 μL of Dynabeads as a pre-clear step. The tube was placed on the magnet and the supernatant removed. This was added to the 50 μL of Dynabeads with the 2C antibody bound and incubated at room temperature shaking for 1 hour. The flow through was removed and added to 2x Laemmli buffer. The beads were washed three times with PBS pH 7.4 with 0.02 % Tween 20 and each wash supernatant retained. Proteins were eluted from the beads by adding 50 μL of 2x Laemmli buffer and heating to 100°C.

## Funding

This work was supported by BBSRC grant (BB/T015748/1) awarded to MRH, DJR and NJS. DMP was funded by a BBSRC DTP studentship (BB/M011151/1). MRH was also supported by the MRC (MR/S007229/1). The funders had no role in the study design, data collection and analysis, decision to publish or preparation of the manuscript.

## Acknowledgements

We thank Francisco Sobrino (Centro De Biologia Molecular Severo Ochoa, Madrid) for the gift of FMDV primary antibodies and Toby Tuthill (The Pirbright Institute, Pirbright) for the gift of purified 3C.

## Author contributions

MRH, NJS and DJR designed the study and wrote the manuscript. DMP conducted the *in vitro* translation experiments. DMP and MRH conducted the replication assays. MRH conducted the immunofluorescence assays. DMP and MRH analysed the data. MRH, NJS and DJR provided supervision.

## Materials & correspondence

Correspondence and materials requests should be directed to MRH.

## Notes

### Competing Interest Statement

The authors have declared no competing interest.

