## Supplemental Figure 1 for "Insights into polyprotein processing and RNA-protein interactions in foot-and-mouth disease virus genome replication"

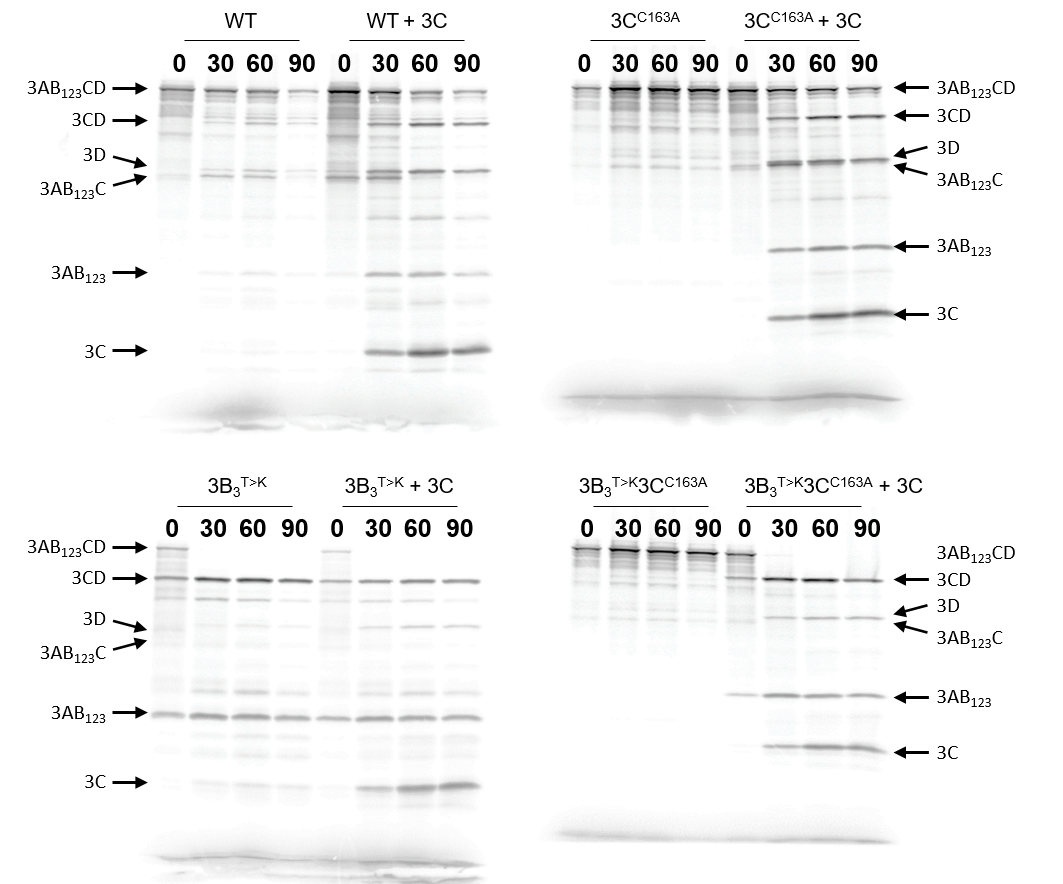

**Figure S1. The 3B_3_^T>K^ point mutation drives *trans*-mediated precursor proteolysis.** Plasmids expressing proteolytically inactive polyproteins with or without the 3B_3_^T>K^ mutation (termed 3B_3_^T>K^3C^C163A^ and 3C^C163A^, respectively) were used to prime coupled [^35^S] labelled transcription/translation assays, followed by unlabelled amino acid chase. To a duplicate set of reactions 10 µM purified 3C^pro^ was added (+3C) immediately after chase. As controls, reactions were setup alongside with the WT or 3B_3_^T>K^ polyproteins without an inactivated 3C^pro^. At regular intervals, samples were taken and reactions stopped by the addition of 2 x Laemmli buffer. Proteins were separated by SDS-PAGE and visualised by autoradiography. The identity of each protein is shown based on predicted molecular weight. Representative gels shown of n=2 experiments.
