## Supplemental Figure 2 for "Insights into polyprotein processing and RNA-protein interactions in foot-and-mouth disease virus genome replication"

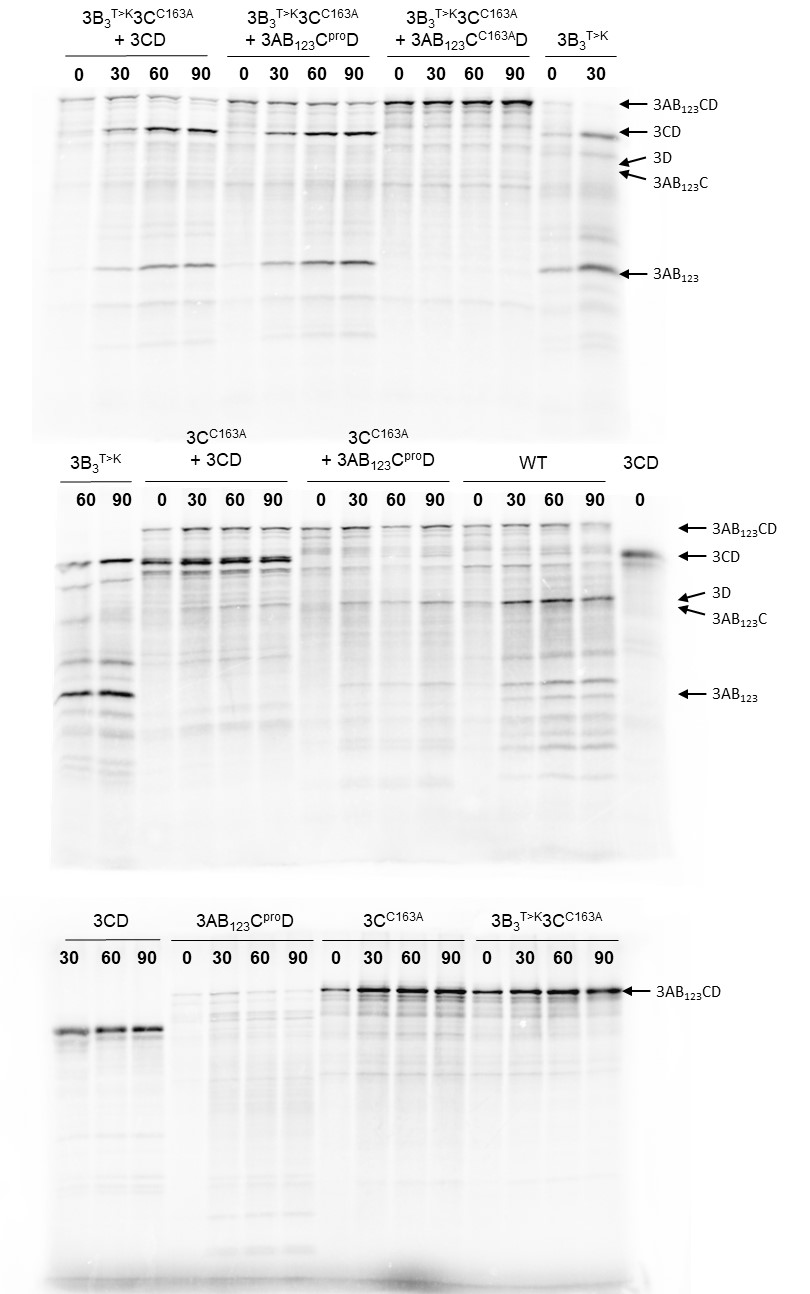

**Figure S2. The 3B_3_^T>K^ point mutation drives *trans*-mediated precursor proteolysis.** Plasmids expressing proteolytically inactive versions of WT or 3B_3_^T>K^ polyproteins (3C^C163A^ and 3B_3_^T>K^3C^C163A^, respectively) were used to prime coupled [^35^S] labelled transcription/translation assays in the presence or absence of 3AB_1,2,3_C^pro^D, a proteolytically active precursor with all cleavage boundaries mutated to prevent self-proteolysis (+3AB_123_C^pro^D). At regular intervals, samples were taken and reactions stopped by the addition of 2 x Laemmli buffer. Proteins were separated by SDS-PAGE and visualised by autoradiography. Representative gels shown of n=2 experiments.
