## Supplemental Figure 4 for "Insights into polyprotein processing and RNA-protein interactions in foot-and-mouth disease virus genome replication"

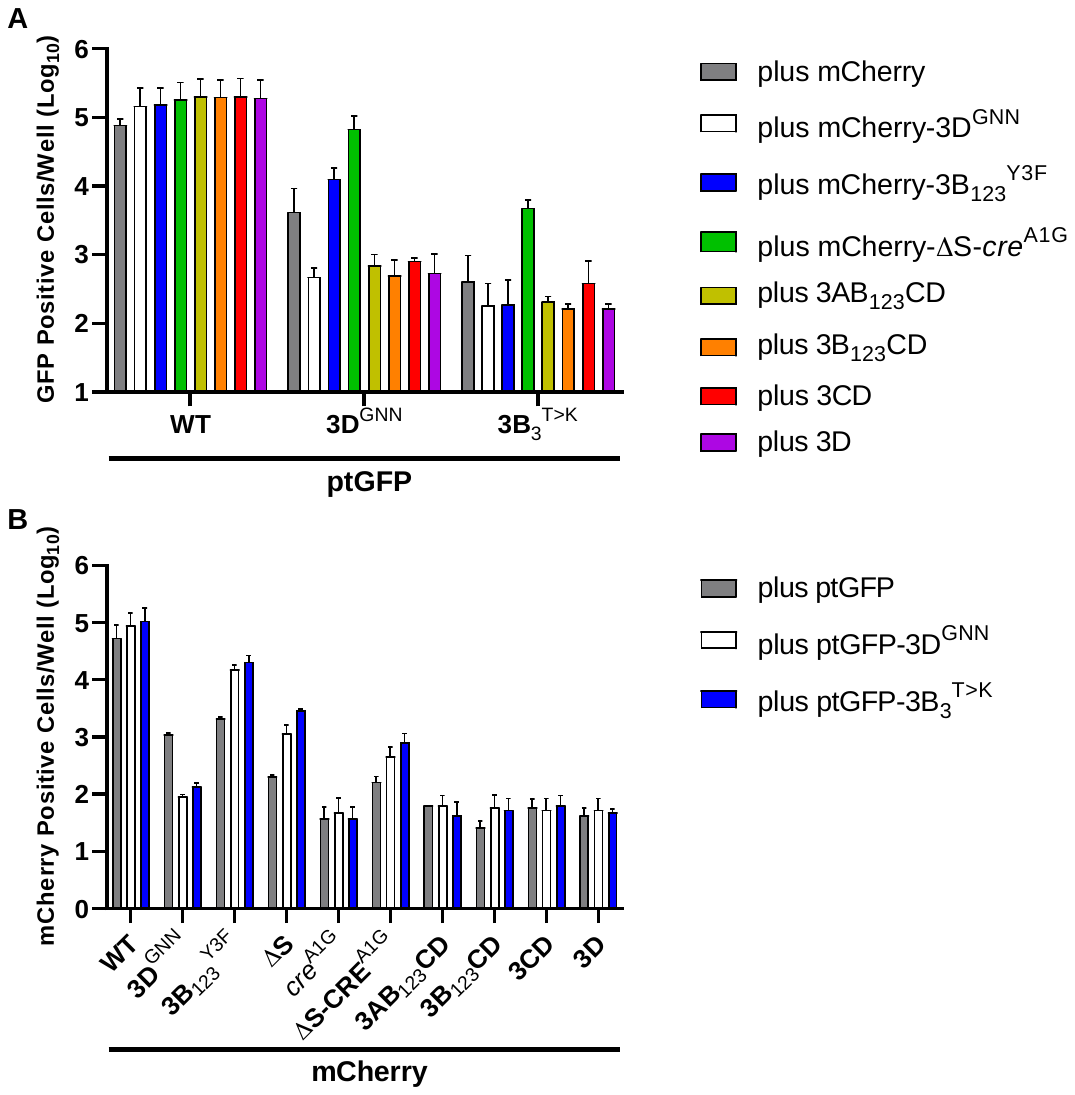

**Figure S4. S-fragment deletions allow *trans*-complementation of *cis-*acting replication components.** BHK-21 cells were co-transfected with mCherry replicons containing S-fragment deletions together with a WT ptGFP, ptGFP-3B_3_^T>K^ or ptGFP-3D^GNN^ replicon. Fluorescent protein expression was monitored hourly for 24 hours. The data show **(A)** ptGFP positive cells per well or **(B)** mCherry positive cells per well at 8 hours post-transfection (n = 2 ± SD).
